# Scale-dependent landscape-biodiversity relationships shape multi-taxa diversity in an oil palm monoculture under restoration

**DOI:** 10.1101/2023.09.08.556058

**Authors:** Denver T. Cayetano, Delphine Clara Zemp, Damayanti Buchori, Sebastian Fiedler, Ingo Grass, Dirk Hölscher, Bambang Irawan, Yevgeniya Korol, Watit Khokthong, Gustavo Brant Paterno, Andrea Polle, Anton Potapov, Leti Sundawati, Teja Tscharntke, Catrin Westphal, Patrick Weigelt, Kerstin Wiegand, Holger Kreft, Nathaly R. Guerrero-Ramírez

## Abstract

Enhancing biodiversity in monoculture-dominated landscapes is a pressing restoration challenge. Tree islands can enhance biodiversity locally, but the role of scale-dependent processes on local biodiversity remains unclear. Using a multi-scale approach, we explored how scale-dependent processes influence the diversity of seven taxa (woody plants, understory arthropods, birds, herbaceous plants and soil bacteria, fauna, and fungi) within 52 experimental tree islands embedded in an oil palm landscape. We show that local, metacommunity (between islands), and landscape properties shaped above- and below-ground taxa diversity, with the stronger effects on above-ground taxa. The spatial extent that best-predicted diversity ranged from 150 m for woody plants to 700 m for understory arthropods with below-ground taxa responding at large spatial extents. Our results underscore the need for multi-scale approaches to restoration. Additionally, our findings contribute to understanding the complex processes shaping multi-taxa diversity and offer insights for targeted conservation and restoration strategies.

## INTRODUCTION

Tropical forests account for at least two-thirds of all terrestrial biodiversity (Gardner *et al*. 2009), one-third of land metabolic activity (Malhi 2012) and offer substantial socio-economic benefits (Foley *et al*. 2005). However, demand for agricultural commodities often results in forest conversion into monoculture-dominated landscapes (Fagan *et al*. 2022; Potapov *et al*. 2022), reducing land-cover diversity critical for taxonomically and functionally diverse communities (Barnes *et al*. 2014; Gámez-Virués *et al*. 2015). With croplands and pastures occupying ∼40% of the land surface (Foley *et al*. 2005), a major challenge is enhancing biodiversity in production areas without compromising yields (Altieri 1999; DeFries *et al*. 2004; Zemp *et al*. 2023).

One strategy for enhancing biodiversity in monoculture-dominated landscapes involves optimizing landscape structure, (i.e., composition & configuration; Foley et al. 2005, Fahrig et al. 2011). Designer landscapes incorporating agroforestry zones can improve ecological connectivity, alleviate edge and matrix effects, and increase landscape heterogeneity (Koh *et al*. 2009; Arroyo-Rodríguez *et al*. 2020). One such implementation, tree islands — clusters of trees integrated into agricultural landscapes through natural regeneration or native tree planting — can enhance local and landscape biodiversity (Corbin & Holl 2012; Montoya-Sánchez *et al*. 2023; Zemp *et al*. 2023) with minimal costs and impediments to yields (Gérard *et al*. 2017; Holl *et al*. 2020; Shaw *et al*. 2020). For example, incorporating tree islands in an oil palm-dominated landscape increased multi-taxa diversity due to changes in vegetation structure and microclimate (Teuscher *et al*. 2016; Zemp *et al*. 2019, 2023; Donfack *et al*. 2021). However, the relative contribution of the individual tree island versus other landscape elements to local biodiversity remains poorly understood.

The unified model of community assembly, an integrated framework of the metacommunity and mainland-island models, may elucidate local biodiversity within tree islands (Figure 1; Fukami 2005, 2009, 2015). The metacommunity model describes interacting communities linked by species dispersal, while the mainland-island model describes species distribution and colonization-extinction dynamics between a “mainland” source habitat and receiving “island” habitat patches (Leibold *et al*. 2004). The unified model considers the interplay of biotic and abiotic factors at various scales. For tree islands within agricultural systems, these broad scales include local properties of the individual tree islands (i.e., *local tree island properties*), interactions among tree islands forming a metacommunity (i.e., *tree island metacommunity*), and the broader landscape context encompassing both tree islands and other land-cover types (i.e., *landscape properties*) (Figure 1).

**Figure 1.**
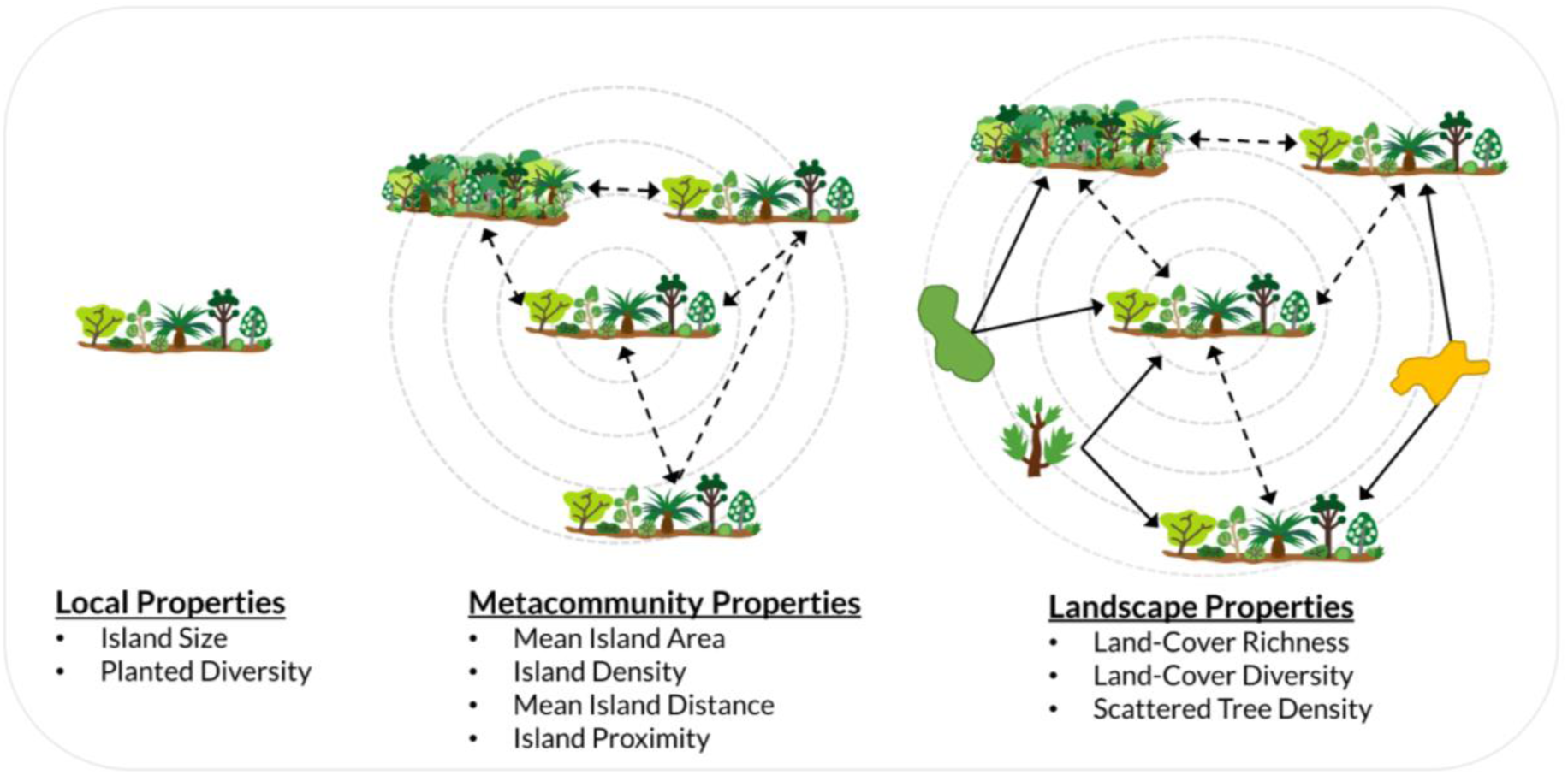
Conceptual illustration of interactions among local, metacommunity, and landscape properties. Under the unified model of community assembly (Fukami 2005, 2009, 2015), multi-taxa diversity within the focal tree island may be jointly determined by these properties. Metacommunity and landscape properties may influence community assembly through internal dispersal among tree islands (dashed arrows), or a combination of internal and external dispersal (solid arrows) from species pools within diverse land-cover types (green and orange polygons) or scattered trees. Dashed circles represent varying spatial extents around the focal tree island. Bullet points denote metrics used as a proxy for each property class. Refer to Table 1 for details on the metrics for each property.

**Table 1.** Tree island, metacommunity, and landscape properties. Metrics describing metacommunity and landscape properties are scale-dependent.

| Type of Metric | Metric name [unit] | Description | Mechanisms |
| --- | --- | --- | --- |
| **Tree island properties (not scale-dependent)** |  |  |  |
| Habitat | Island size [m<sup>2</sup>] | Area of the focal tree island | Larger island size leads to greater community diversity due to the species-area relationship and island biogeographic mechanisms. |
| Habitat | Planted diversity [n] | The number of planted tree species in the focal tree island. | Planted diversity promotes community diversity through varied habitats, resources, and ecological niches, facilitating the coexistence of species with diverse requirements and positive interactions. |
| **Metacommunity properties (scale-dependent)** |  |  |  |
| Area and edge | Mean island area<br>[ha] | The mean area of tree islands in a given spatial extent excluding the focal patch.<br>Usually called “mean patch area.” | Larger mean island areas provide more available habitat and resources, allowing for the colonization and persistence of more species. |
| Aggregation | Island density<br>[islands/100 ha] | Number of tree islands per 100 ha, excluding the focal tree island. | Greater island density increases opportunities for colonization and species interactions, leading to higher species richness and diversity. |
| Aggregation | Mean island distance [m] | Mean Euclidean distance between the focal tree island and each neighboring tree island in meters. | Lower mean island distance enhances species dispersal, connectivity, and interactions, increasing species richness and overall diversity. |
| Isolation | Island proximity<br>[unitless] | The sum of each island area divided by its squared Euclidean distance to the focal tree island (i.e., proximity index). | Greater island proximity enhances species diversity by facilitating dispersal and gene flow, thereby reducing the effects of isolation in a monoculture landscape. |
| **Landscape properties (scale-dependent)** |  |  |  |
| Diversity and source | land-cover richness<br>[land-cover types/100 ha] | The number of land-cover types in the landscape (max = 8, Table S1) per 100 ha.<br>Includes tree islands as a land-cover type.<br>Does not consider the relative abundance of land-cover types. | Greater land-cover richness provides a higher density of patches with diverse species assemblages, supporting greater interactions, unique ecological roles, and niches. |
| Diversity and source | land-cover diversity<br>[unitless] | Shannon's index based on the proportion of landscape occupied by each land-cover type (Verheyen *et al.* 1999; McGarigal *et al.* 2012)), excluding focal tree island (Table S1). | Greater land-cover diversity provides varied habitats, resources, and ecological niches, enhancing species richness and facilitating species movement for increased diversity. |
| Source | Scattered tree density [trees /100 ha] | The number of scattered trees per 100 ha (Korol *et al.* 2021). | Scattered tree density provides seeds, microhabitats, resources, and connectivity, supporting a greater variety of species. |

In the context of tree islands, *local tree island properties* such as island size and planted diversity influence what species can colonize and survive (Teuscher *et al*. 2016; Zemp *et al*. 2023). On a larger scale, the *tree island metacommunity* may support biodiversity through habitat heterogeneity, facilitation of rescue effects and source-sink dynamics (Leibold *et al*. 2004; Blitzer *et al*. 2012; Borthagaray *et al*. 2015). In addition, the broader *landscape properties* encompass other land-cover types that may include keystone structures acting as biodiversity sources and refuges (Duelli & Obrist 2003; Tews *et al*. 2004; Theron *et al*. 2020; van der Mescht *et al*. 2023). Landscape structure can influence the availability of habitats, and routes and barriers for species dispersal (Reid *et al*. 2014; Martin *et al*. 2019). Therefore, community assembly in tree islands may be driven by combinations of local, metacommunity, and landscape properties influencing various ecological responses, including species interactions and dispersal (Fahrig et al. 2011, Miguet et al. 2016). However, their effect on multi-taxa diversity in the tropics and their relevance to biodiversity enhancement in monoculture-dominated landscapes remain uncertain.

Interactions between components of landscape structure (i.e., metacommunity and landscape properties) and biodiversity tend to vary with spatial scales and taxa (Wiens 1989; Holland *et al*. 2004; Jackson & Fahrig 2015; Newman *et al*. 2019). Therefore, understanding landscape-biodiversity interactions requires consideration of the “scale of effect” — the spatial extent that best predicts population response after evaluating these interactions across multiple spatial extents (Brennan *et al*. 2002; Jackson & Fahrig 2012). This scale reflects species mobility or dispersal ability, which influences how different taxa interact with the landscape (Jackson & Fahrig 2012; Miguet *et al*. 2016). However, the complexity of these relationships often limits studies to few species and spatial extents (Jackson & Fahrig 2015). Consequently, there is a need for parsimonious approaches to integrate multi-taxa landscape-biodiversity relationships into landscape design. Investigations into the underlying mechanisms shaping biodiversity patterns, including between-patch dispersal, and landscape diversity and connectivity, are seldom carried out in restoration contexts. Thus, filling this gap could inform targeted landscape restoration strategies to enhance biodiversity.

In this study, we look through the lens of the unified model of community assembly to examine the drivers of local diversity in seven above- and below-ground taxa in tree islands within an oil palm-dominated landscape. Specifically:

**1.** We investigated the influence of local, metacommunity, and landscape properties on tree island biodiversity across different taxa and spatial extents. We hypothesized that metacommunity properties enhance local taxa diversity by improving landscape connectivity and that landscape properties such as secondary forests and scattered trees also contribute by serving as species sources (Le Provost *et al*. 2021). Further, we expected that the properties and spatial extent that best predict biodiversity may be associated with body size which is related to overall taxa mobility (see Miguet et al. 2016). Specifically, properties at smaller spatial extents (including local properties) may primarily affect less mobile taxa such as soil fauna. In contrast, highly mobile taxa such as birds may be influenced mainly by larger-scale metacommunity and landscape properties.
**2.** We determined the most parsimonious modeling approach to inform the design of restoration landscapes that support multi-taxa diversity by comparing univariate and multivariate models. We hypothesized that multivariate models would best predict local taxa diversity by capturing processes acting at local, metacommunity, and landscape scales. Therefore, by selecting key properties at different scales, we can provide parsimonious strategies for landscape design that consider multi-taxa diversity.

## MATERIALS AND METHODS

### Study Site

Our study site is an oil palm-dominated landscape near Bungku village in the Jambi province of Sumatra, Indonesia (01.95° S & 103.25° E; 47 ± 11 m a.s.l.; Figure 2). Mean rainfall in this region is 2235 ± 385 mm yr^-1^, and mean temperature is 26.7 ± 1.0 °C yr^-1^. The soil type is predominantly loamy Acrisol (Allen *et al*. 2015), and the pre-conversion forest was dipterocarp-dominated lowland rainforest (Drescher *et al*. 2016; Teuscher *et al*. 2016). The EFForTS Biodiversity Enrichment Experiment (EFForTS-BEE; www.uni-goettingen.de/crc990) was established in December 2013 and includes 52 tree islands of varying sizes (25, 100, 400 & 1600 m^2^) and planted tree richness (0, 1, 2, 3, 6 species; hereafter “planted diversity,” Table 1) distributed across an oil-palm dominated landscape covering 140 ha (Zemp *et al*. 2023). Planted trees were selected from a pool of six native, multi-use species (see Teuscher et al. (2016) and Zemp et al. 2019a for details on the experimental design). As of 2016, the landscape was dominated by oil palm (82%), with minor coverage of fallow land (6%), roads and bare soil (5% each), secondary forests (4%), rubber plantations (2%), and a combined 1% for orchards, urban areas, and water bodies (Figure 2; Khokthong 2019).

**Figure 2.**
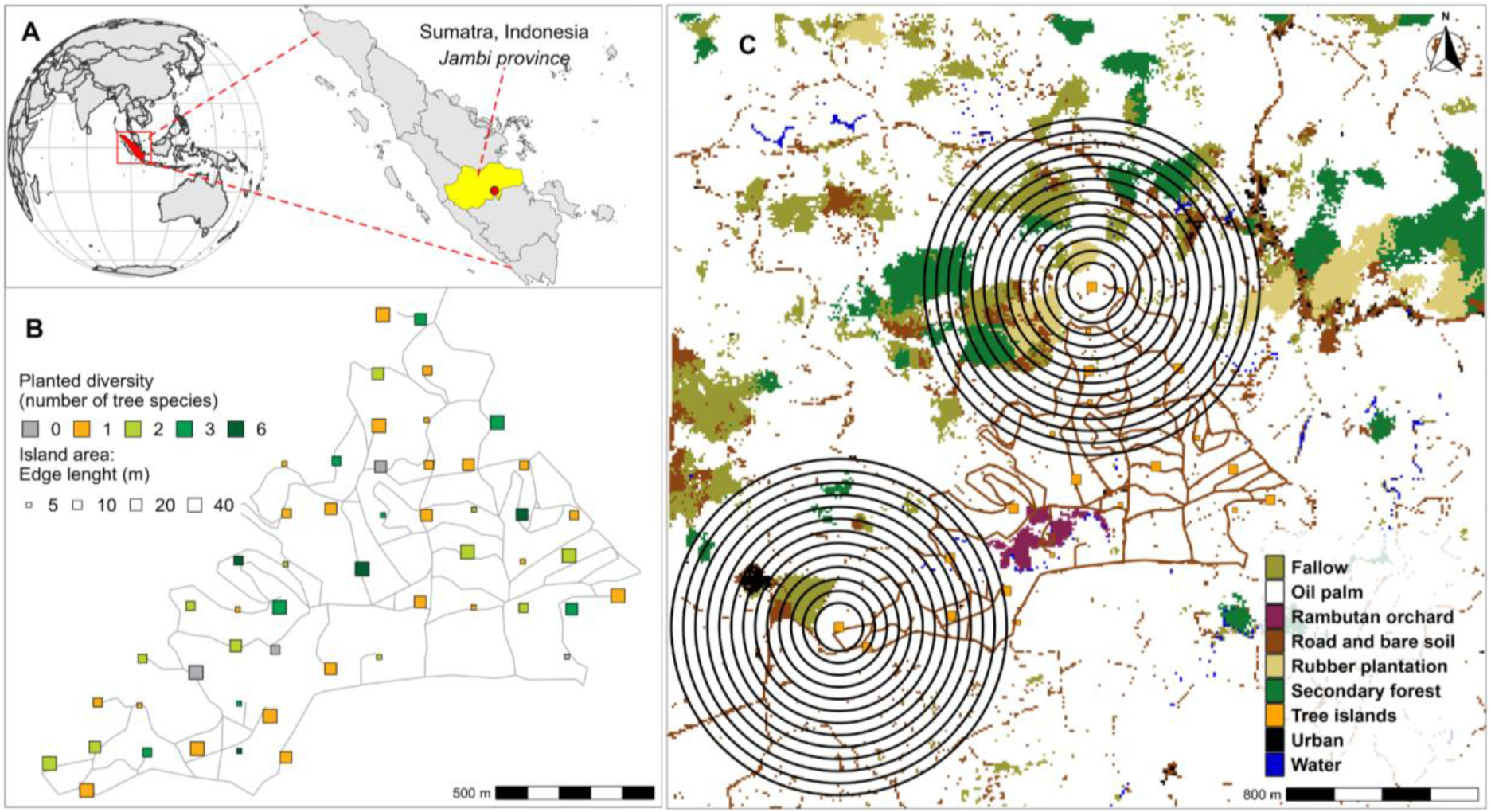
Map of the study site showing A.) the geographical location of EFForTS-BEE; B.) tree island spatial distribution, sizes, and planted diversity; C.) exemplary layout of the evaluated spatial extents (max = 700 m) around two tree islands.

### Assessing Above- and Below-Ground Taxa Diversity

We surveyed the diversity of seven taxa — birds, woody plants, herbaceous plants, understory arthropods, soil fauna, soil fungi, and bacteria — of each tree island between October 2016 and August 2018 (Zemp *et al*. 2023). The sampling approach varied by taxon: **birds** were sampled visually and with sound recordings from four 15-minute point counts within 28 m of the tree island center (Darras *et al*. 2018). Free-standing **woody plants** (shrubs, trees, and bamboo) with a height of ≥ 1.3 m were censused within each tree island. As the sampling area for woody plants corresponded to tree island area, we standardized diversity estimates using rarefaction curves to 24 individuals, the median number of individuals per plot (Zemp *et al*. 2023). **Herbaceous plants** were identified to genus, species, or morphospecies levels and consisted of all non-woody vascular plants of < 1.3 m in height (excluding epiphytes) within a randomly placed 25 m^2^ subplot in each tree island. **Understory arthropods** were sampled three times using two sets of three adjoining pan traps filled with water and a drop of detergent. Each pan trap was colored with yellow ultraviolet paint to attract pollinators (Westphal *et al*. 2008) and placed at the height of understory plants with an exposure time of 45 hours. **Soil fauna** were sampled from soil collected from four distinct 256 cm^2^ areas (5 cm depth including litter) within each 25 m^2^ subplot. All invertebrates were extracted with Kempson heat extractors (Kempson *et al*. 1963) and identified to family/order level (30 groups in total; Potapov et al. 2019). **Soil fungi** were sampled from three soil cores (10 cm depth, 4 cm diameter), excluding roots and litter layer. The fungal community was assessed using Illumina next-generation sequencing (Illumina Inc., San Diego, USA) and further classified into operational taxonomic units (OTU) using the BLAST algorithm (BlastN, v2.7.1; Camacho et al. 2009) and the UNITE v7.2 (UNITE_public_01.12.2017.fasta; Kõljalg et al. 2013). See Ballauff et al. (2020) for details on sampling, sequencing, and classification. **Soil bacteria** were sampled from three soil cores (10 cm) 1 m away from adjacent trees. DNA and RNA extraction and posterior classification were then conducted on the homogenized and root-free samples (Berkelmann *et al*. 2020). See Zemp et al. (2023) for more information on biodiversity sampling.

For each tree island, we calculated diversity using the effective number of species q=1 (Jost 2006, Chao et al. 2014). Compared to other diversity metrics, the effective number of species q=1 neither favors locally rare (species richness, q=0) nor abundant species (Simpson diversity, q=2; Roswell et al. 2021).

### Local, Metacommunity, and Landscape Properties

We selected seven metrics to characterize local, metacommunity, and landscape properties (Table 1, Figure 1). Selected metrics included two metrics associated with *local tree island properties* (island size & planted diversity), four metrics associated with *tree island metacommunity properties (*mean island area, island density, mean island distance, & island proximity), and three associated with *landscape properties* (land-cover richness, land-cover diversity, & scattered tree density; see Table 1 for metric descriptions, Figure 1).

Metacommunity and landscape metrics were calculated using a 10 m resolution land-cover map (3.3 × 3.3 km) with eight land-cover classes mapped in 2016 using red-blue-green and near-infrared imagery taken from a fixed-wing drone. Land-cover types were determined by supervised classification with the maximum likelihood classifier (Figure 2 & Tables S1 & S2, see Khokthong (2019) for details on image capture and processing). Scattered tree density was calculated based on a map of 10,652 scattered trees across the landscape (10.1 trees ha^-1^) identified from high-resolution drone imagery (3.3 × 3.3 km). See Korol et al. (2021) for details on scattered tree identification and mapping.

We calculated each metric (Table 1) at twelve different spatial extents centered around each focal tree island following the focal patch multi-scale approach (Brennan et al. 2002). Spatial extents ranged from radii of 150 m to 700 m with increments of 50 m to capture the scale of effect of taxa with weak dispersal ability. The lower limit was based on the mean nearest neighbor distance between tree islands to ensure each spatial extent included more than the focal tree island (mean = 140.2 m, min = 105.6 m, max = 182 m). The upper limit was determined by the tree island nearest to the map boundary, ensuring an equal number of scales across all tree islands. If any land-cover patches or tree islands intersected the boundary of the spatial extent, only portions within the boundary were considered in calculations. Focal tree islands were excluded from metric calculations to separate tree island properties as fixed effects in the models. All metrics were calculated in R version 4.1.0 (R Core Team 2021) using the tidyverse (Wickham *et al*. 2019) and sf (Pebesma 2018) packages and following specifications outlined by FRAGSTATS (McGarigal *et al*. 2012; McGarigal 2015).

### Statistical Analyses

Statistical analyses were conducted using R version 4.1.0 (R Core Team 2021). Metrics were transformed to ensure normality of residuals using the “bestNormalize” function from the “bestNormalize package” (version 1.8.2; Figure S1; Peterson and Cavanaugh 2020). Other R packages used included tidyverse (Wickham *et al*. 2019), broom (Robinson *et al*. 2021), and sf (Pebesma 2018).

First, we evaluated correlations between metacommunity and landscape metrics (Figures S2 & S3). We used univariate linear regressions at each spatial extent to evaluate scale-dependent effects of each metric on taxa diversity. Scale-dependent variation was visualized by plotting the standardized model coefficients against spatial extent. Scale-dependency was determined by 1) identifying the scale of effect — the spatial extent with the largest coefficient estimate — and the spatial extent with the lowest estimate, and 2) comparing these two models using the Akaike information criterion (AIC). Relationships where the difference in AIC (ΔAIC) was greater than 2, indicating improved model fit, were considered scale-dependent as per Jackson and Fahrig (2015).

To understand the complementary effects and relative importance of local, metacommunity, and landscape properties on multi-taxa diversity, we performed multivariate models including all metrics for each taxon at each spatial extent. We subsequently performed model selection for each taxon per spatial extent. Specifically, we ran all possible model combinations, identified the model with the lowest AIC (i.e., the best model) and models with ΔAIC <= 2 from the best model, and performed model averaging (full average) with the identified models, using the “dredge” and “mod.avg” functions of the MuMIn package (version 1.46.0; Barton 2020).

We assessed spatial autocorrelation of model residuals using the global Moran’s I index, using the “lm.morantest” function from the spdep package (version 1.2-8; Bivand 2022). After applying a Bonferroni correction to account for multiple testing, spatial autocorrelation was found statistically significant in 14% of univariate and 13% of multivariate models (P < 0.05). However, Moran’s I index values were relatively low (0.04 - 0.15 and -0.06 - 0.11, respectively), indicating weak spatial autocorrelation. Local Moran’s I analysis on the diversity estimates for each taxon and metric across scales revealed similarly weak autocorrelation (Figures S4 & S5). Therefore, no further correction was implemented.

## RESULTS

### Local Multi-Taxa Diversity

Local diversity across tree islands varied between 2.81 ± 1.85 for birds, 7.26 ± 2.99 for herbaceous plants, 3.41 ± 2.2 for woody plants (excluding planted trees), 23.55 ± 6.69 for understory arthropods, 3456.99 ± 958.23 for soil bacteria, 174.19 ± 49.33 for soil fungi and 6.16 ± 1.29 for soil fauna.

### Variation in Metacommunity and Landscape Properties Across Scales

Metacommunity metrics either increased, decreased, or remained constant across spatial extents (Figure S6). Specifically, mean island distance and island proximity increased with spatial extent, with mean island distance ranging from 120.47 ± 40.99 to 409.40 ± 31.90 m and island proximity from 0.05 ± 0.06 to 0.18 ± 0.07 at 150 and 700 m radius, respectively. In contrast, island density decreased with spatial extent, varying from 26.66 ± 16.25 to 16.89 ± 4.31 hm^-1^ while mean island area remained relatively consistent across spatial extents ranging from 0.03 ± 0.04 to 0.05 ± 0.01 ha. Landscape metrics increased with spatial extent, with land-cover richness ranging from 61.49 ± 13.69 to 5.55 ± 0.42, land-cover diversity from 0.33 ± 0.17 to 0.61 ± 0.20, and scattered tree density increasing from 176.02 ± 246.22 to 554.38 ± 217.42 hm^-1^ (Figure S6).

### Scale-Dependent Metacommunity- and Landscape-Taxa Diversity Relationships

Metacommunity- and landscape-taxa diversity relationships showed scale-dependency as indicated by ΔAIC > 2 between models with the largest (i.e., scale of effect) and the smallest estimates (63% of cases, Figure S7). Scale dependency was mostly associated with changes in the strength instead of the direction of the relationship, except for the relationship between island density and bird diversity (Figure 3). In this case, an increase in island density was associated with lower bird diversity at the smallest spatial extent, while at larger spatial extents (> 500 m), the inverse pattern was observed. Island proximity was the most consistent predictor across spatial extents and taxa.

**Figure 3.**
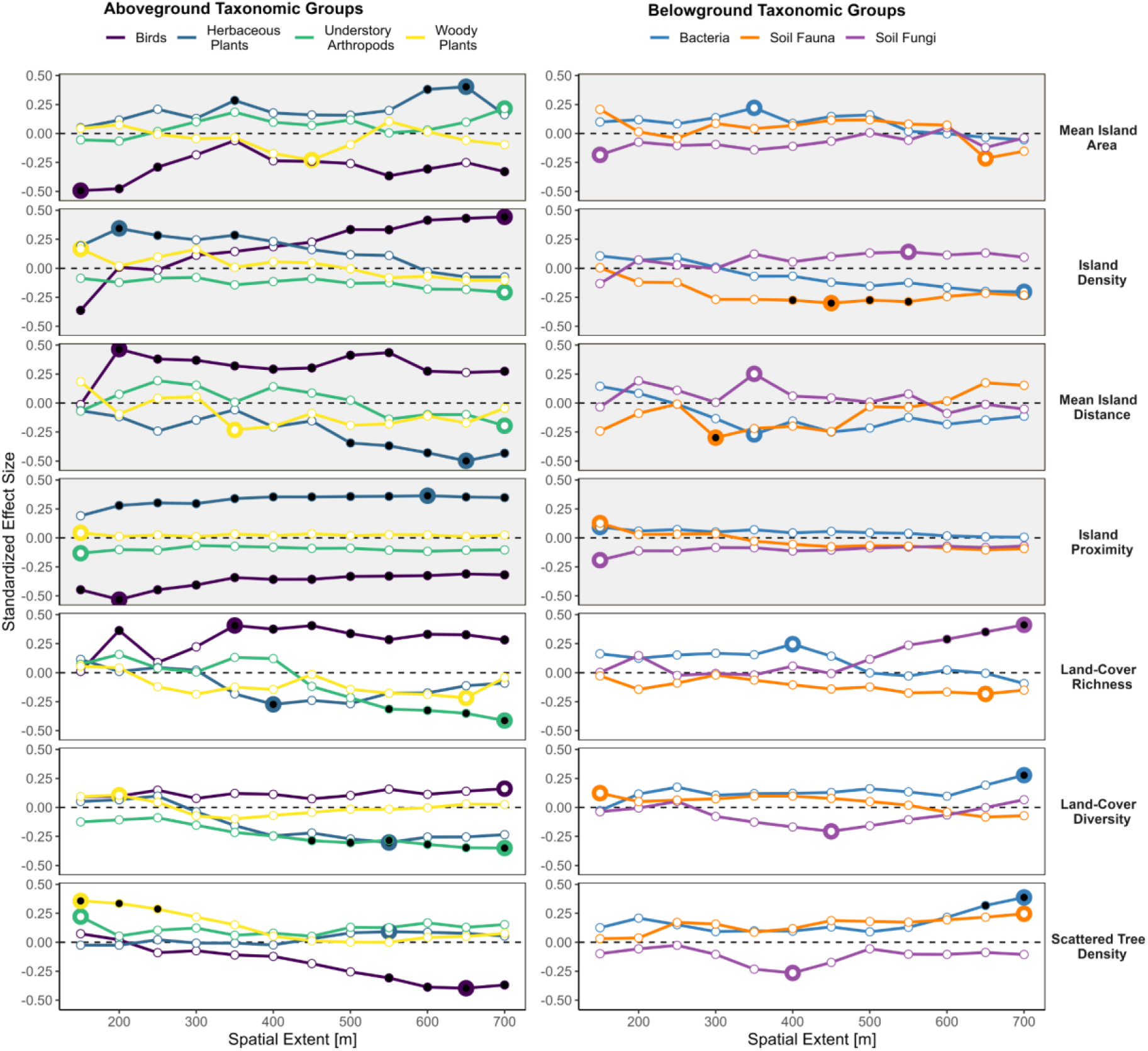
Above- and below-ground taxa-diversity responses to metacommunity (grey panels) and landscape properties (white panels). Standardized effect sizes are taken from the univariate linear models. Spatial extents are shown as radii from the center of the focal tree island in meters. Large circles show the scale of effect — the largest coefficient estimate. Black-filled circles show significant effects (P <=0.05).

Metacommunity and landscape properties influenced above-ground taxa more strongly than below-ground taxa diversity across spatial extents (Figure 3). For above-ground taxa, changes in local diversity of birds and herbaceous plants were mostly associated with metacommunity properties. However, for these specific taxa, the scale of effect — the scale with the largest coefficient estimate — and the direction of the relationship with metacommunity properties (except island density) followed a contrary trend. For instance, bird diversity was mainly associated with metrics at small scales (< 250 m), while herbaceous plant diversity was associated with larger scales (> 550 m). Furthermore, bird diversity decreased with higher mean island area (scale of effect: 150 m, β = -0.49, P < 0.001) and island proximity (scale of effect: 200 m, β = -0.53, P < 0.001) and increased, overall, with higher island density (scale of effect: 700 m, β = 0.44, P = 0.001) and mean island distance (scale of effect: 200 m, β = 0.47, P = 0.001). In contrast, herbaceous plant diversity increased with higher mean island area (scale of effect: 650 m, β = 0.40, P = 0.003), island density (scale of effect: 200 m, β = 0.34, P = 0.013), and island proximity (scale of effect: 600 m, β = 0.28 to 0.36, P < 0.05) and decreased with higher mean island distance (scale of effect: 650 m, β = -0.50, P < 0.001). For below-ground taxa, changes in local diversity of soil fauna were associated with metacommunity properties, with soil fauna diversity decreasing with higher island density (scale of effect: 450 m, β = -0.30, P = 0.031) and mean island distance (scale of effect: 300 m, β = -0.30, P = 0.031).

Idiosyncratic patterns for the scale of effect and direction of landscape-taxon diversity relationships were mainly observed for above-ground taxa (Figure 3). Specifically, bird diversity increased with land-cover richness (scale of effect: 350 m, β = 0.41, P = 0.003) and decreased with scattered tree density (scale of effect: 650 m, β = -0.40, P = 0.004). Herbaceous plants and understory arthropods diversities decreased with land-cover richness (scale of effect: 400 m, β = -0.27, P = 0.05 and 700 m, β = -0.41, P = 0.002, respectively) and land-cover diversity (scale of effect: 550 m, β = -0.30, P = 0.029 and 700 m, β = -0.35, P = 0.011, respectively). Woody plant diversity increased with scattered tree density (scale of effect: 150 m, β = 0.36, P = 0.009). In contrast to the above-ground taxa, the below-ground taxa responded to the landscape properties with a consistent scale of effect of 700 m. Here, soil fungi diversity increased with higher land-cover richness (β = 0.41, P = 0.002), while soil bacteria diversity increased with higher land-cover diversity (β = 0.28, P = 0.046) and scattered tree density (β = 0.39, P = 0.005).

### Univariate Versus Multivariate Models for Modelling Taxa Responses

To inform parsimonious approaches for designing biodiversity-friendly landscapes, we compared the effectiveness of univariate versus multivariate models involving local, metacommunity, and landscape properties in capturing taxa diversity. Our results show that multivariate models considerably enhanced variance explained for most of the taxa at the scale of effect, except for bacteria and fungi (Figure 4). In the case of bacteria, the multivariate model only marginally improved the explained variance compared to the univariate model (Adjusted R^2^ = 0.16 and R^2^ = 0.15, respectively). For soil fungi, the highest variance was explained by the univariate model with land-cover richness (R^2^ = 0.17) than in the best multivariate model (adjusted R^2^ = 0.14). The multivariate models explained higher variance than the best univariate model for all other taxa. For example, the variance explained by the best univariate model for bird diversity (i.e., island proximity at 200 m) was 0.28, increasing to 0.37 in the best multivariate model (550 m; Figure 4). Further, the variance explained increased from 0.25 to 0.30 for herbaceous plants, from 0.13 to 0.40 for woody plants, from 0.90 to 0.27 for soil fauna, and from 0.17 to 0.36 for understory arthropods.

**Figure 4.**
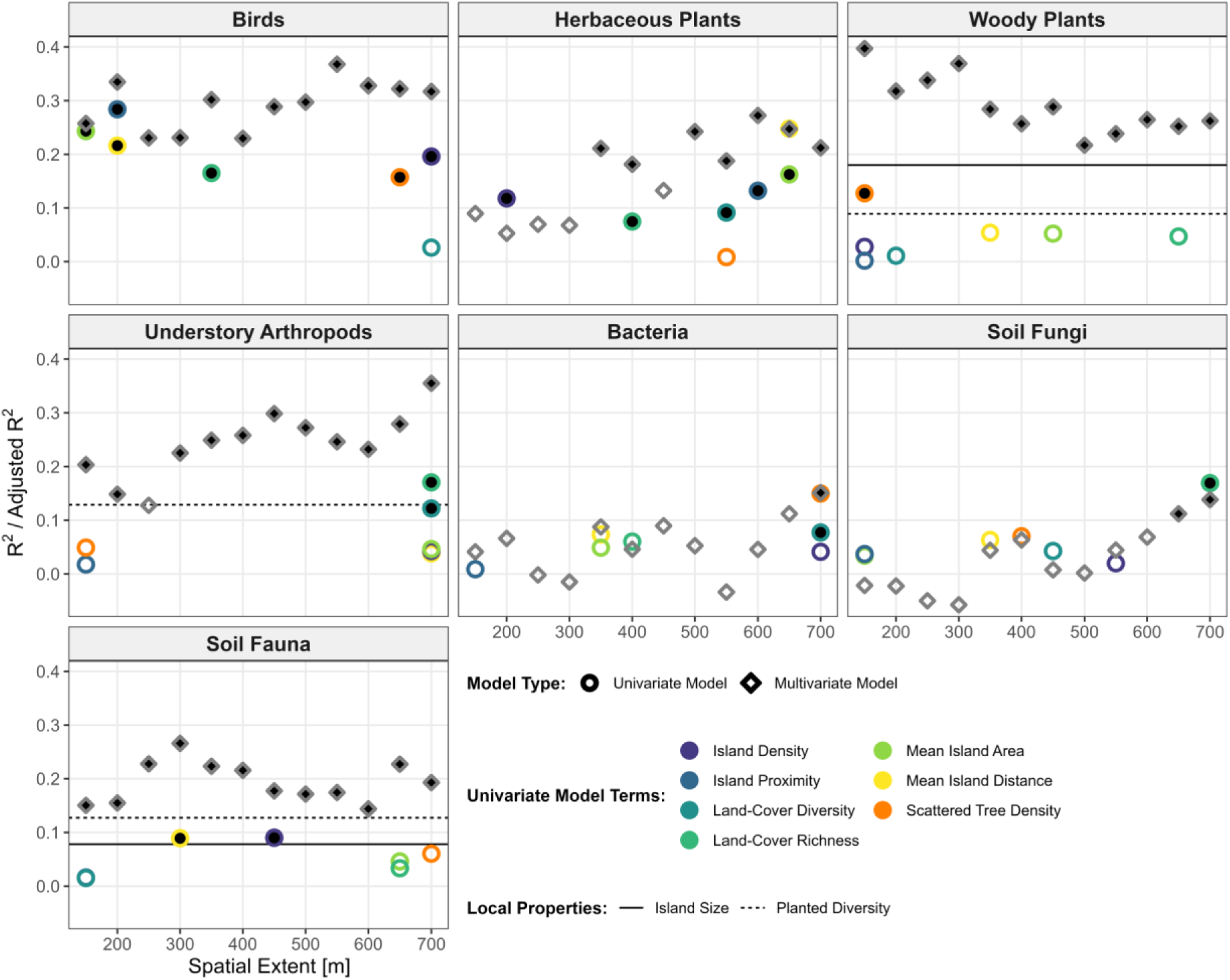
Explanatory power of univariate (circles) and multivariate models (diamonds) displayed using R^2^ and adjusted R^2^, respectively. Univariate models are shown at the scale of effect, while multivariate models span all spatial extents. Multivariate models incorporate all metrics (including tree island size and planted diversity) and are the best after model selection and averaging. Filled symbols indicate significance (P <0.05). Solid/dashed lines represent significant (P < 0.05) relationships between taxa diversity and local properties. Spatial extents are shown as radii from the center of the focal tree island in meters.

### Relative Contribution of Local, Metacommunity, and Landscape Properties to Local Multi-Taxa Diversity

The best multivariate models combined local, metacommunity, and landscape properties across all taxa except soil fauna. Notably, metacommunity properties were consistently included in the best multivariate models across taxa, except birds and herbaceous plants, but with confidence intervals that often overlapped zero suggesting weak or variable responses. The multivariate scales of effect varied by taxa: 150 m for woody plants, 300 m for soil fauna, 550 m for birds, 600 m for herbaceous plants, and 700 m for soil fungi, bacteria, and understory arthropods (see Figures 4 & 5).

Woody plant diversity was mainly influenced by combinations of local and landscape properties, specifically island size (β = 0.42) and scattered tree density (β = 0.39). Soil fauna diversity was primarily associated with island size (β = 0.31) and planted diversity (β = -0.35), both local properties. Metrics associated with metacommunity, or landscape properties were simultaneously included in the best-averaged models across taxa. Bird diversity was shaped by a combination of local and metacommunity properties, encompassing island size (β = 0.34), island density (β = 0.29), and mean island distance (β = 0.42). Herbaceous plants were driven mainly by metacommunity properties such as mean island area (β = 0.35) and mean island distance (β = -0.34). Soil fungi and bacteria diversity were most influenced by landscape properties, specifically land-cover richness (fungi: β = 0.46) and scattered trees (β = 0.37), respectively. Understory arthropods revealed a combined influence of local and landscape properties, with planted diversity (β = -0.34) and land-cover richness (β = -0.34) both having strong negative effects.

**Figure 5.**
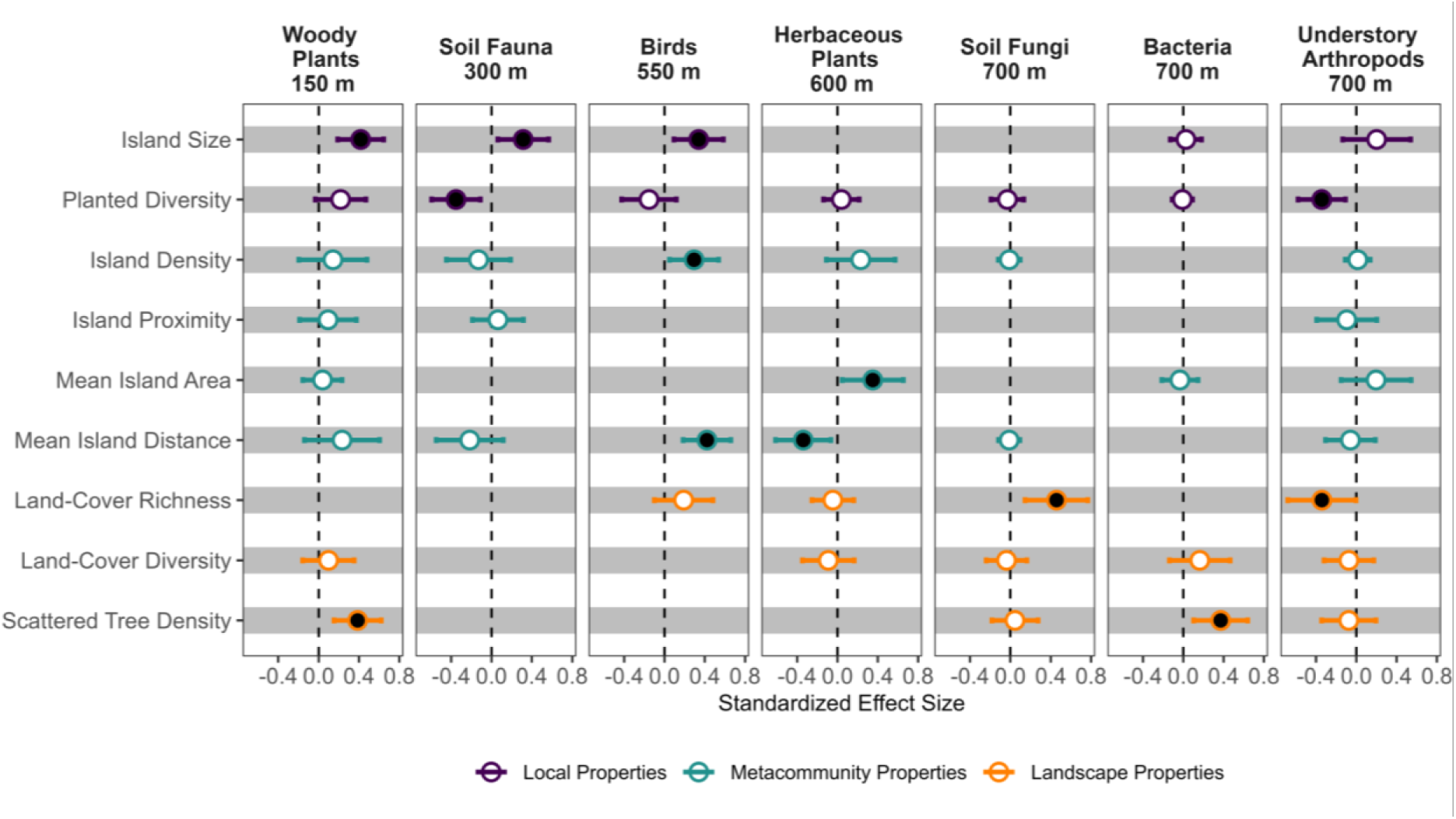
The best multivariate models at the scale of effect for each taxon. The panel headers identify the taxa and scales of effect. Filled symbols indicate confidence intervals not overlapping zero. Effect sizes from the multivariate models are standardized. Best-averaged models across spatial extents are shown in Figure S8.

## DISCUSSION

We drew upon the unified model of community assembly to provide insights into enhancing multi-taxa diversity in tropical monoculture-dominated landscapes. We identified distinct influences of broad spatial properties on biodiversity. For instance, tree island metacommunity properties primarily shaped bird and herbaceous plant diversity, while landscape properties influenced soil fungi, bacteria, woody plants, and understory arthropod diversity. Relationships between the landscape components (i.e., local, metacommunity, & landscape properties) and taxa diversity suggest the nature of these relationships may hinge on variations in mobility, dispersal strategies, and life history among taxa. Further, we show that relationships between biodiversity and both metacommunity and landscape properties are largely scale-dependent, supporting general expectations (Wiens 1989; Jackson & Fahrig 2012, 2015). However, contrary to the expected size-mobility linkages (Stevens *et al*. 2014; Hillaert *et al*. 2018), neither small-bodied (e.g., arthropods & soil fauna) nor large-bodied taxa (e.g., birds) showed a distinct preference for small or large spatial extents. Interestingly, below-ground taxa diversity was related to both local properties (e.g., soil fauna), and landscape properties (e.g., bacteria & soil fungi). This challenges the longstanding view that below-ground communities are solely influenced by local conditions, a shift observed in temperate (Le Provost *et al*. 2021), and now in tropical landscapes. Finally, by integrating local, metacommunity, and landscape properties into our models, we improved their ability to explain variations in taxa diversity. This highlights the importance of a holistic strategy to enhance biodiversity in monoculture-dominated landscapes, considering processes across spatial scales, and explicitly considering landscape context.

### Above- and Below-Ground Responses to Local, Metacommunity, and Landscape Properties

Above- and below-ground communities exhibited different responses to local, metacommunity, and landscape properties, highlighting the need to extend the focus of landscape ecology from plants, birds, and insects (see review in Jackson and Fahrig 2015) to below-ground taxa (Mennicken et al. 2020). Below-ground responses are often assumed to be influenced by processes and properties acting at local scales (Tscharntke *et al*. 2012). However, this perspective oversimplifies the complexity and diversity of below-ground taxa. In the best multivariate model, soil fauna was indeed mainly shaped by local conditions. However, soil fungi and bacteria responded strongly to landscape properties, specifically to land-cover richness and scattered tree density at larger spatial extents. The influence of landscape properties at large spatial extents on below-ground diversity has also been observed in temperate human-modified ecosystems (Le Provost *et al*. 2021), suggesting this pattern may span biomes. It is likely that despite dispersal limitations for soil fungi and bacteria (Grilli *et al*. 2017; Mony *et al*. 2022), land-cover types such as fallow, secondary forests, and orchards, as well as scattered trees, act as species sources. Specifically, different land-cover types and scattered trees may increase landscape heterogeneity and serve as habitats for diverse communities shaped by variations in soil abiotic conditions, microhabitats, microclimates, light conditions, and resource availability, among others (Grilli *et al*. 2017; Mony *et al*. 2022). Furthermore, more mobile above-ground taxa may assist in transporting below-ground taxa and/or their resources across the landscape (Moore *et al*. 1988); however, this remains highly speculative.

Tree island age may influence the observed effects of local and metacommunity properties on taxa diversity. In their early stages, taxa requiring longer maturation periods (e.g, trees), or those with delayed responses such as below-ground communities (Mennicken *et al*. 2020), may depend on land-cover types other than tree islands and scattered trees as species sources. As they mature, the influence of metacommunity properties is expected to increase with species reaching reproductive ages and establishing populations. Further, island age may also influence local biotic or abiotic properties, modifying the influences of environmental filtering on community assembly. Therefore, a temporal perspective is necessary considering the multidecadal nature of the restoration process (Holl *et al*. 2017; Guerrero-Ramírez 2021) and how effects change over time as the succession occurs. Our results suggest that environmental filters act at local and landscape scales, likely determining observed taxa responses to landscape properties and spatial extent through influences on generalist-specialist dynamics. Landscapes may moderate diversity patterns by selecting functionally relevant traits, thereby reducing functional redundancy and response diversity following habitat modification (Tscharntke *et al*. 2012). Forest specialists are more affected by the loss of primary habitat than generalists, who can use resources from a wide range of habitats (Pardini *et al*. 2010; Galán-Acedo *et al*. 2018; Morante-Filho *et al*. 2018).

For example, birds evaluated in this study exhibited a wide range of diets (i.e., granivores, insectivores, and omnivores) and habitat preferences (i.e., wooded & cultivated areas, forest gaps & edges, primary & secondary forests) but over 96% showed a preference for agricultural or disturbed habitats (Figure S9). Since secondary forests comprise only 4% of the study area, environmental filtering might have already impacted the species composition (Figure 1). This is particularly significant given the recommendation to maintain at least 40% forest in a landscape to conserve both forest species and habitat generalists (Arroyo-Rodríguez *et al*. 2020). Therefore, taxa responses to landscape properties and spatial extent should be interpreted in the context of ecological traits of species in the landscape.

The influence of metacommunity and landscape properties on bird diversity are likely related to high dispersal abilities, which enable easy movement through the anthropogenic matrix, especially at smaller spatial extents (Knowlton *et al*. 2017). Specifically, higher bird diversity is associated with increased density of small but distant tree islands, according to the multivariate models. These findings suggest these islands play a crucial role in facilitating movement by enhancing habitat connectivity. Additionally, small tree islands can increase ecotone lengths and habitat interspersion/juxtaposition, leading to greater resource availability and foraging efficiency of birds, especially for omnivores and/or habitat generalists (Morante-Filho *et al*. 2018).

Herbaceous plant diversity was influenced by metacommunity properties, increasing with mean island area, island density, and proximity but decreasing with mean island distance. Interestingly, this pattern is inverse to that observed in birds. These contrasting responses underscore the importance of considering taxon-specific responses for conservation and restoration, and understanding how biotic interactions shape ecological processes across scales. For instance, decreased plant diversity could result from bird-dispersed invasive plants dominating herbaceous communities (Sachsenmaier 2018), further reflecting the implications of a shift towards omnivore-dominant bird populations (Figure S5).

The negative relationship between land-cover richness and arthropod diversity observed in our study suggests that elements beyond land-cover richness may influence the direction of this relationship. For example, the size of the regional species pool, patch quality, and degree of specialization (with specialists less likely to disperse to tree islands) could drive community assembly in the tree islands. Fewer land-cover types with higher similarity to tree islands, such as secondary forests and orchards, may act as more important species sources than other types, such as rubber plantations. This expectation is supported by a positive relationship between the similarity index and arthropod diversity (Figure S3). Arthropod preference for specific land-cover types could be related to resource availability, plant structural complexity, and phenology, among others (Stamps & Linit 1998).

Additionally, simplified plant communities, such as the oil palm matrix, can reduce insect diversity and promote single-species proliferation, often as pests (Risch *et al*. 1983; Stamps & Linit 1998; Brandmeier *et al*. 2021). This may explain contrasting relationships between land-cover richness and arthropod diversity observed in temperate zones, where increased land-cover richness has been associated with higher arthropod diversity (Gámez-Virués *et al*. 2015).

## CONCLUSION

Our research reveals scale-dependent responses of multiple taxa during the restoration process in a monoculture-dominated landscape. We show the importance of not only individual tree islands but also the tree island metacommunity and other landscape properties in enhancing biodiversity. We observe that above-ground taxa were generally influenced more by metacommunity and landscape properties, whereas local or landscape properties primarily drove below-ground taxa. This accentuates the necessity of considering these scale-dependent properties for the preservation of not only above-ground but also below-ground taxa, supporting a shift already advocated for in temperate regions but not tested in the tropics. Notably, these taxon-dependent responses might be linked to ecological traits such as mobility, dispersal, and life history. For example, metacommunity properties greatly impacted taxa with high mobility and dispersal, like birds and herbaceous plants, while landscape properties were crucial for taxa with overall lower dispersal, such as understory arthropods and tree species. These patterns might fluctuate with island maturity, further emphasizing the importance of preserving landscape attributes such as forests and scattered trees. This underlines that creating and managing tree islands as a restoration strategy should take a landscape perspective informed by multi-scale analyses.

## ACKNOWLEDGEMENTS

This study was funded by the Deutsche Forschungsgemeinschaft (DFG, German Research Foundation) – project ID 192626868 – SFB 990 in the framework of the collaborative German-Indonesian research project CRC990. We thank PT Humusindo for granting us access to their properties. D. T. C. acknowledges the German Academic Exchange Service (DAAD) for providing financial support through its “Development-Related Postgraduate Courses” (EPOS) funding program. WK received a Ph.D. fellowship from the Royal Government of Thailand within the Development and Promotion of Science and Technology Talents Project (DPST). N. G-R thanks the Dorothea Schlözer Postdoctoral Programme of the Georg-August-Universität Göttingen and the financial support of Deutsche Forschungsgemeinschaft (DFG), grant number 316045089/GRK 2300. CW is grateful for being funded by the DFG (project number 493487387).

