## Supporting Information for "Scale-dependent landscape-biodiversity relationships shape multi-taxa diversity in an oil palm monoculture under restoration"

### Supplementary Methods

Metacommunity and landscape properties

Initially, we selected nine metrics to characterize metacommunity and landscape properties, including island coverage and the similarity index. However, due to high correlations between these two metrics and others, both island coverage and the similarity index were excluded from the analysis (Figures S2 and S3). Island coverage, measured in percentage, represents the fraction of the total landscape area (i.e., spatial extent) occupied by tree islands, influencing connectivity between habitats and shaping community dynamics. The similarity index, expressed as a value between 0 and 1, measures the similarity between different land-cover types, reflecting landscape homogeneity or heterogeneity. This can impact ecological processes like species colonization and competition and indicate ecosystem resilience to disturbances. To calculate the similarity index, similarity coefficients for each land-cover type were determined based on expert knowledge (Table S3).

### Supporting Tables

**Table S1.** Definition of land-cover classes. Modified from Khokthong (2019).

| Land Cover Class | Definition |
|---|---|
| Secondary forest | Patches of forest and disturbed forest, which are bigger than 0.5 ha. From the prior classification maps in 2013, the forest area in Bungku village and the study site was indicated as the secondary forest. |
| Oil palm plantation | Cultivated area for palm-oil agriculture. Young oil palms were defined as oil palm plantations that should have a canopy size of at least 1 m in the same homogeneous field. |
| Rubber plantation | The agricultural land used for planting rubbers and partially mixed with tree species. Some other tree species' canopy could be seen where rubbers' canopy is dominant. |
| Rambutan orchard | Rambutan plantation |
| Fallow | Predominant grasses or shrubs where natural vegetation growth may take place after disturbance. |
| Water | Streams, canals, and non-flowing water as lakes. |
| Urban | Area of intensive houses in the village. Individual household structures. The residential area and construction site are included in this class. |
| Bare soil | Barren soil without tree cover or pathway for transportation. |

**Table S2.** Confusion matrix displaying the accuracy assessment of the land-cover map derived from Khokthong (2019)**.**

| | | Reference data | | | | | | | | | |
|---|---|---|---|---|---|---|---|---|---|---|---|
| | **Land use types in 2016** | Secondary forest | Oil palm plantation | Rubber plantation | Orchard | Fallow | Water | Urban | Bare soil | Total | User's accuracy (%) |
| **Classification data** | Secondary forest | **51** | 4 | 0 | 0 | 8 | 0 | 0 | 0 | 63 | 80.95 |
| | Oil palm plantation | 0 | **58** | 0 | 0 | 2 | 0 | 0 | 3 | 63 | 92.06 |
| | Rubber plantation | 0 | 3 | **54** | 0 | 6 | 0 | 0 | 0 | 63 | 85.71 |
| | Orchard | 6 | 5 | 0 | **41** | 1 | 1 | 0 | 9 | 63 | 65.08 |
| | Fallow | 7 | 8 | 1 | 0 | **39** | 4 | 0 | 4 | 63 | 61.90 |
| | Water | 1 | 0 | 0 | 1 | 1 | **58** | 0 | 2 | 63 | 92.06 |
| | Urban | 0 | 1 | 0 | 0 | 0 | 1 | **59** | 2 | 63 | 93.65 |
| | Bare soil | 0 | 4 | 1 | 0 | 4 | 0 | 0 | **54** | 63 | 85.71 |
| | Total | 65 | 83 | 56 | 42 | 61 | 64 | 59 | 74 | **504** | |
| | Producer's accuracy (%) | 78.46 | 69.88 | 96.43 | 97.62 | 63.93 | 90.63 | 100 | 72.97 | | |
| | **Overall accuracy (%)** | **82.14** | | | | | | | | | |

**Table S3.** Similarity coefficients assigned to various land cover types or classes, used in calculating the similarity index for the study. The coefficients represent the degree of similarity for each land cover type, ranging from 1 for high similarity (e.g., tree islands, secondary forest) to 0.1 for low similarity (e.g., urban areas, roads & bare soil).

| | **Land Cover Type** | **Similarity Coefficient** |
|---|---|---|
| 1 | Tree islands | 1 |
| 2 | Secondary forest | 1 |
| 3 | Rubber Plantation | 0.7 |
| 4 | Rambutan Orchard | 0.7 |
| 5 | Fallow | 0.4 |
| 6 | Oil Palm | 0.2 |
| 7 | Water | 0.1 |
| 8 | Urban | 0.1 |
| 9 | Road & Bare Soil | 0.1 |

### Supporting Figures

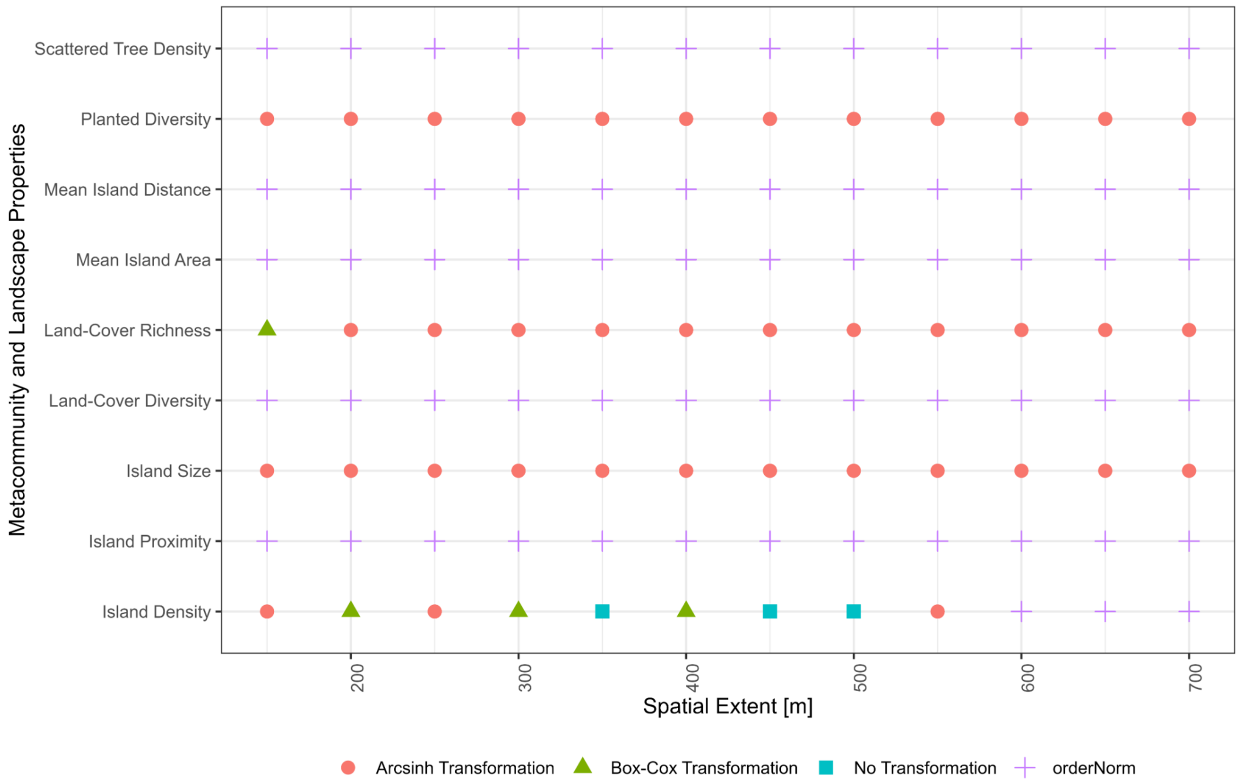


**Figure S1.** Transformation type used from the *bestnormalize* R package (version 1.8.2; Peterson & Cavanaugh, 2020) to transform the local, metacommunity, and landscape properties to a normal distribution.

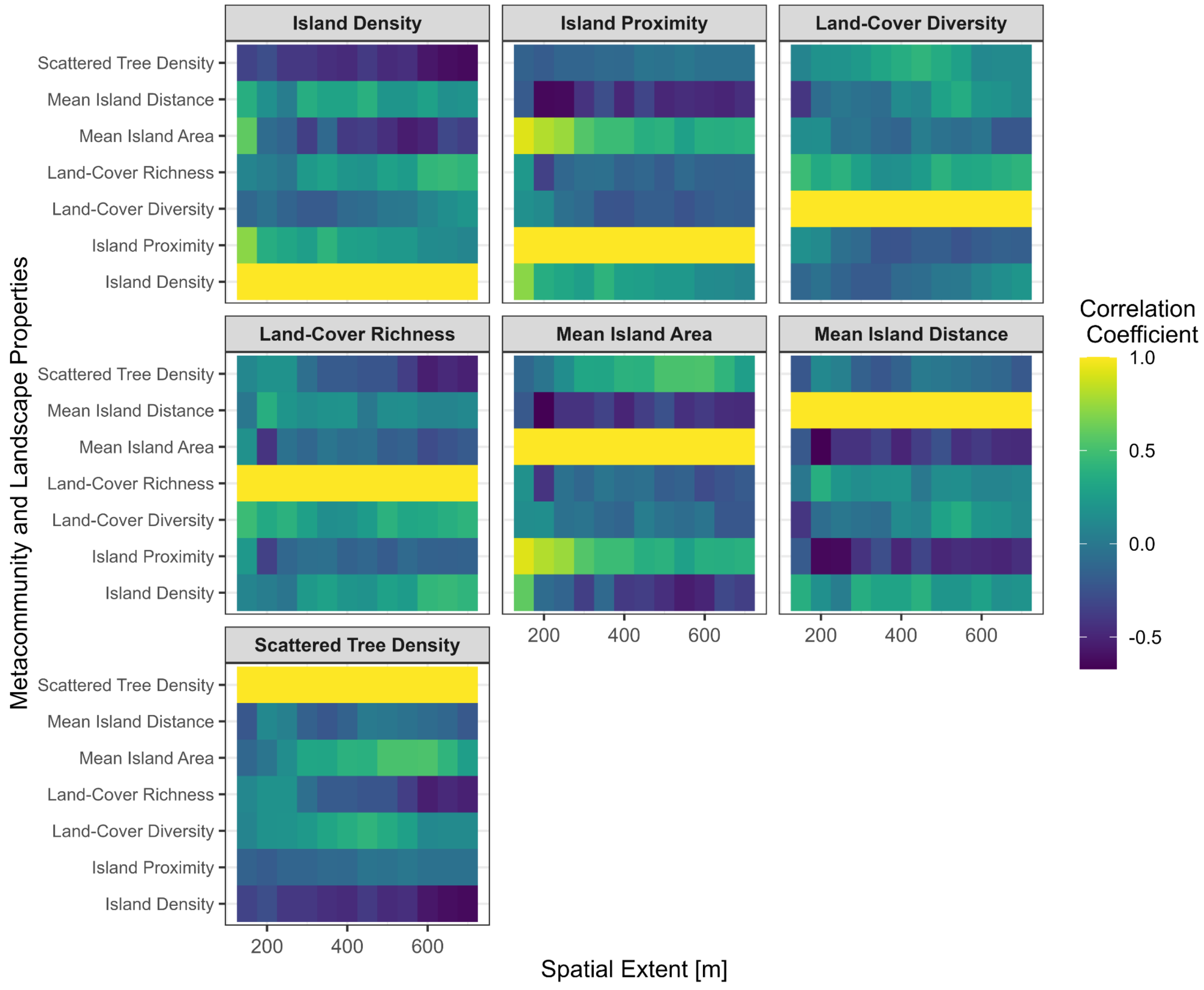


**Figure S2.** Pearson's Correlations of Metacommunity and Landscape Metrics Across Spatial Extents. This figure delineates the interrelationships between tree island metacommunity properties (mean island area, island density, mean island distance, island proximity, island coverage) and landscape properties (land-cover richness, land-cover diversity, scattered tree density, land-cover similarity) across various spatial scales. The correlation coefficients are calculated using Pearson's method. The color gradient in the plot represents the strength and direction of the correlations.

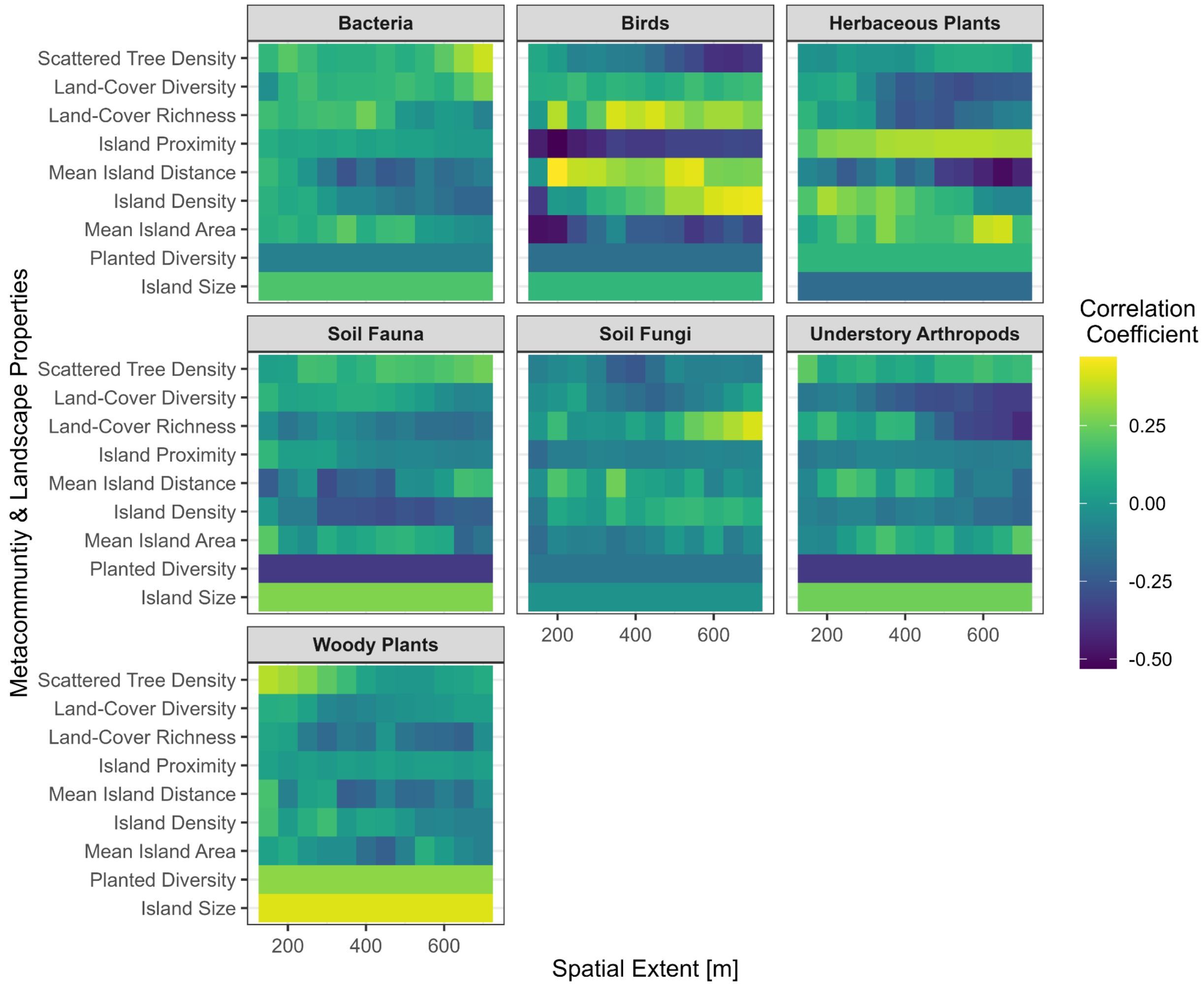


**Figure S3.** Pearson's Correlation between local, metacommunity, and landscape properties across spatial extents. This figure explores the correlations between local (island size and planted diversity), metacommunity (mean island area, island density, mean island distance, island proximity, island coverage), and landscape metrics (land-cover richness, land-cover diversity, scattered tree density, land-cover similarity). The correlation analysis is conducted using Pearson's method. The color gradient in the plot represents the strength and direction of the correlations.

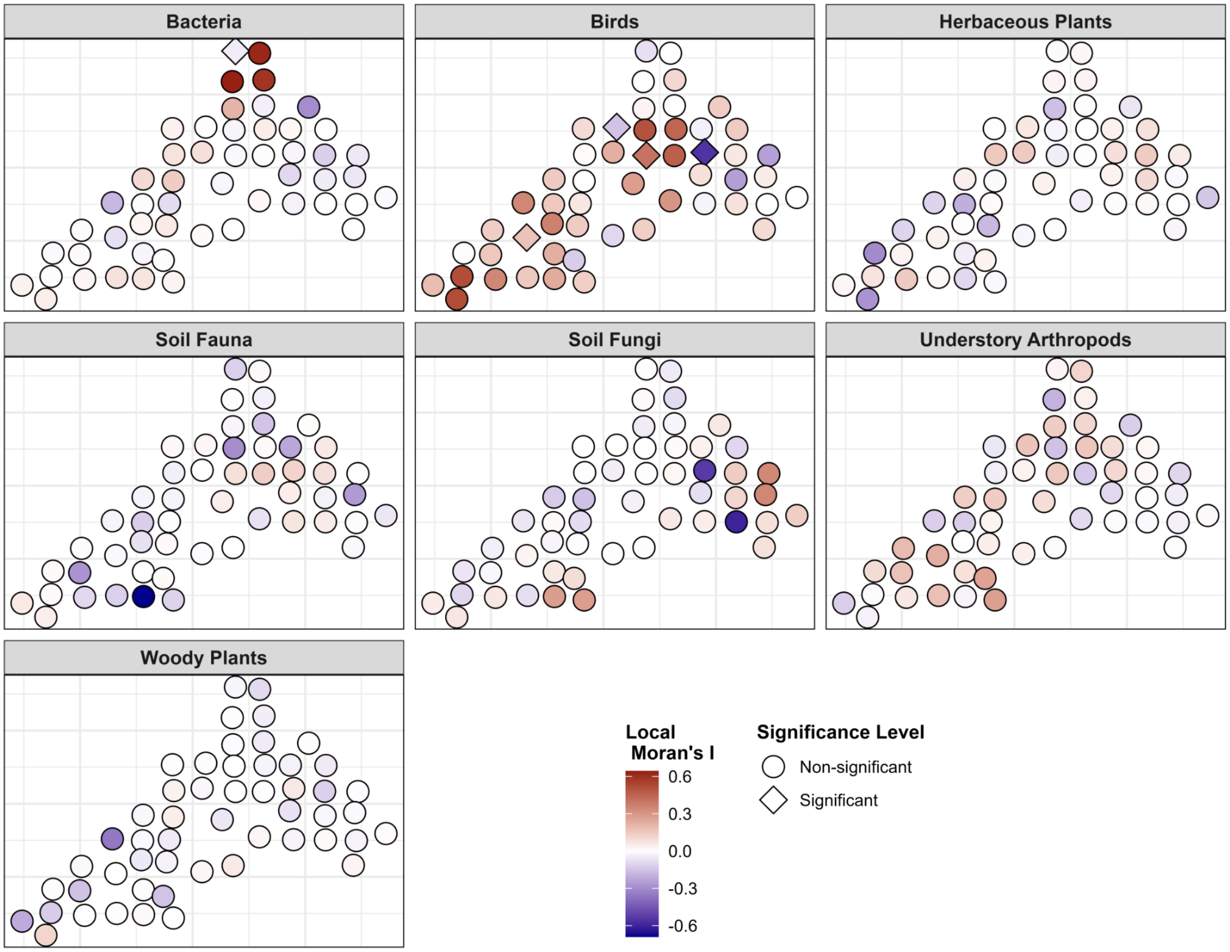


**Figure S4.** Local Moran's statistic for spatial autocorrelation of taxa diversity within tree islands. This figure employs the Local Moran's statistic to measure spatial autocorrelation between taxa diversity within individual tree islands. Tree islands with significant spatial autocorrelation (p-value <= 0.05) are highlighted with a diamond symbol. The color gradient in the plot represents the strength and direction of the spatial autocorrelation.

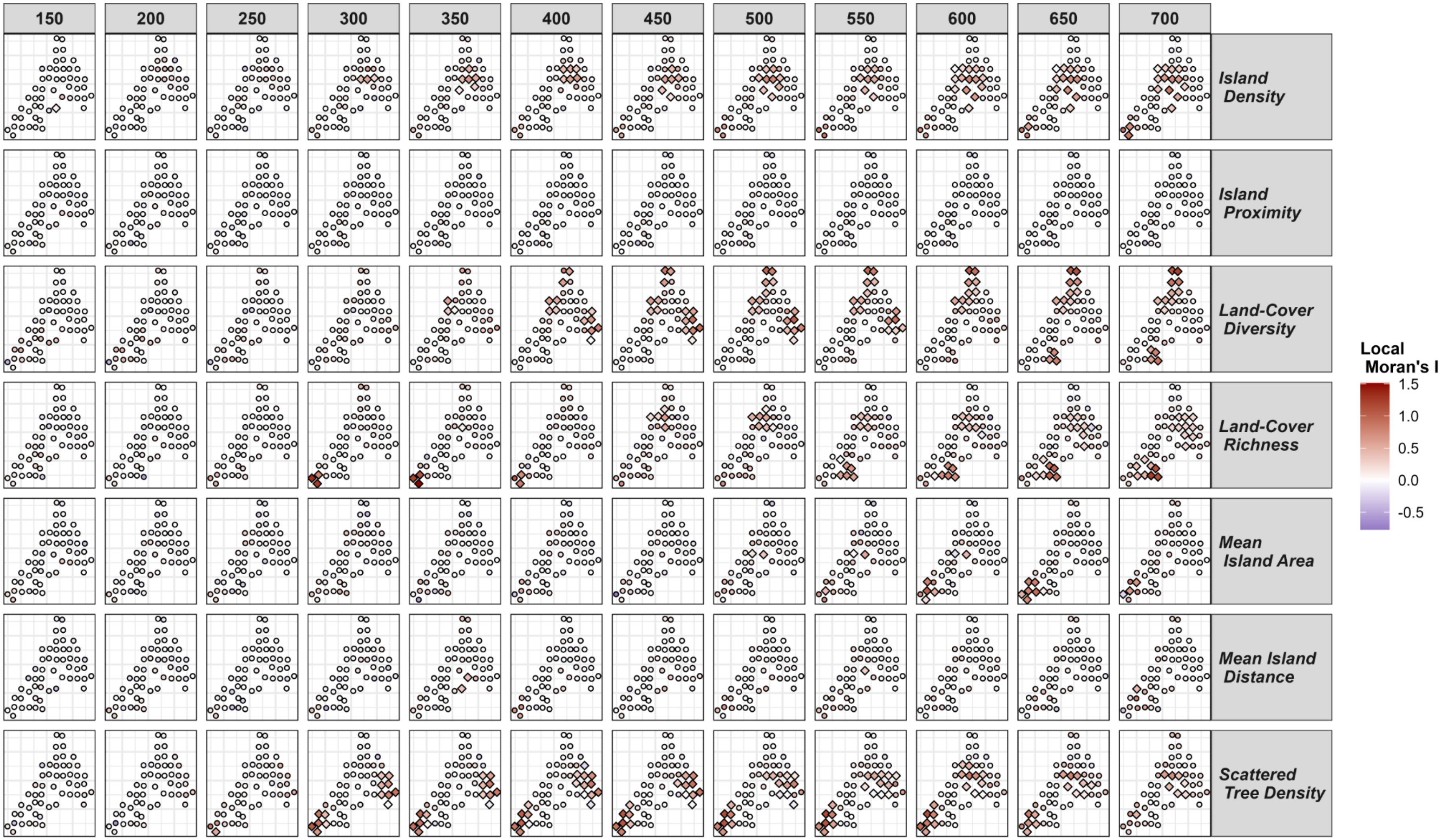


**Figure S5.** Local Moran statistic measures spatial autocorrelation between landscape metrics across spatial extents. Tree islands with significant values (p-value <=0.05) are shown with a triangle symbol with the local Moran statistic above.

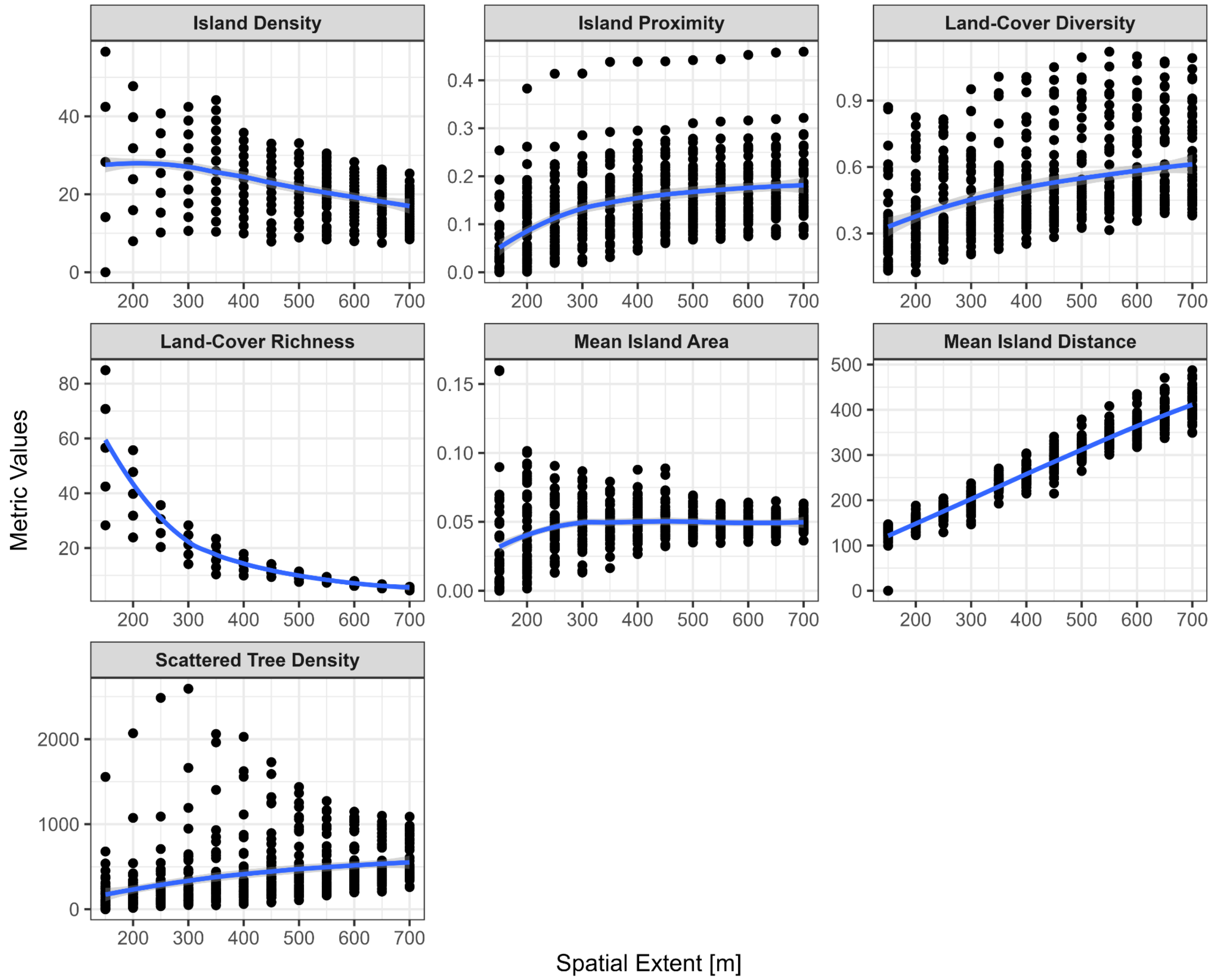


**Figure S6.** Landscape metrics as a function of spatial extent. Panels show a.) mean island area; b) mean island distance; c) island density; d) island percentage; e) island proximity; f) land-cover richness; g) land-cover diversity; h) similarity index; i) scattered tree density. Spatial extent is represented by the radius from the center of its focal tree island in meters. LOESS curve is shown in blue. Units of the metrics can be found in Table 1. Points represent values for each tree island (n =52 at each spatial extent).

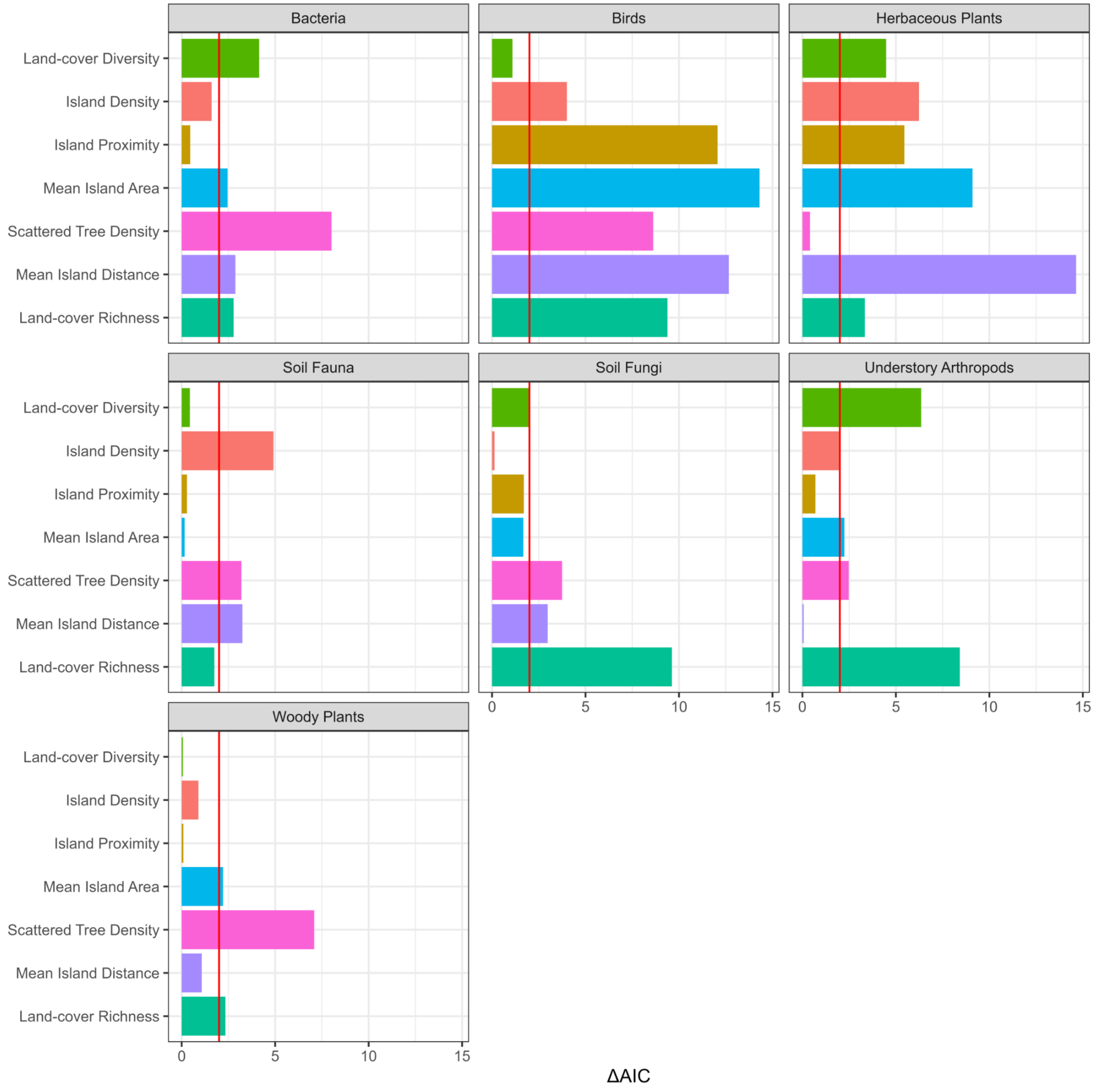


**Figure S7**. Differences in AIC (ΔAIC) between the smallest and largest (i.e., the scale of effect) effect sizes across spatial extents for various taxa-metric relationships. The graph shows that the ΔAIC was greater than two in 63% of the cases, a threshold represented by the red horizontal line. Each bar color represents different metacommunity or landscape metrics.

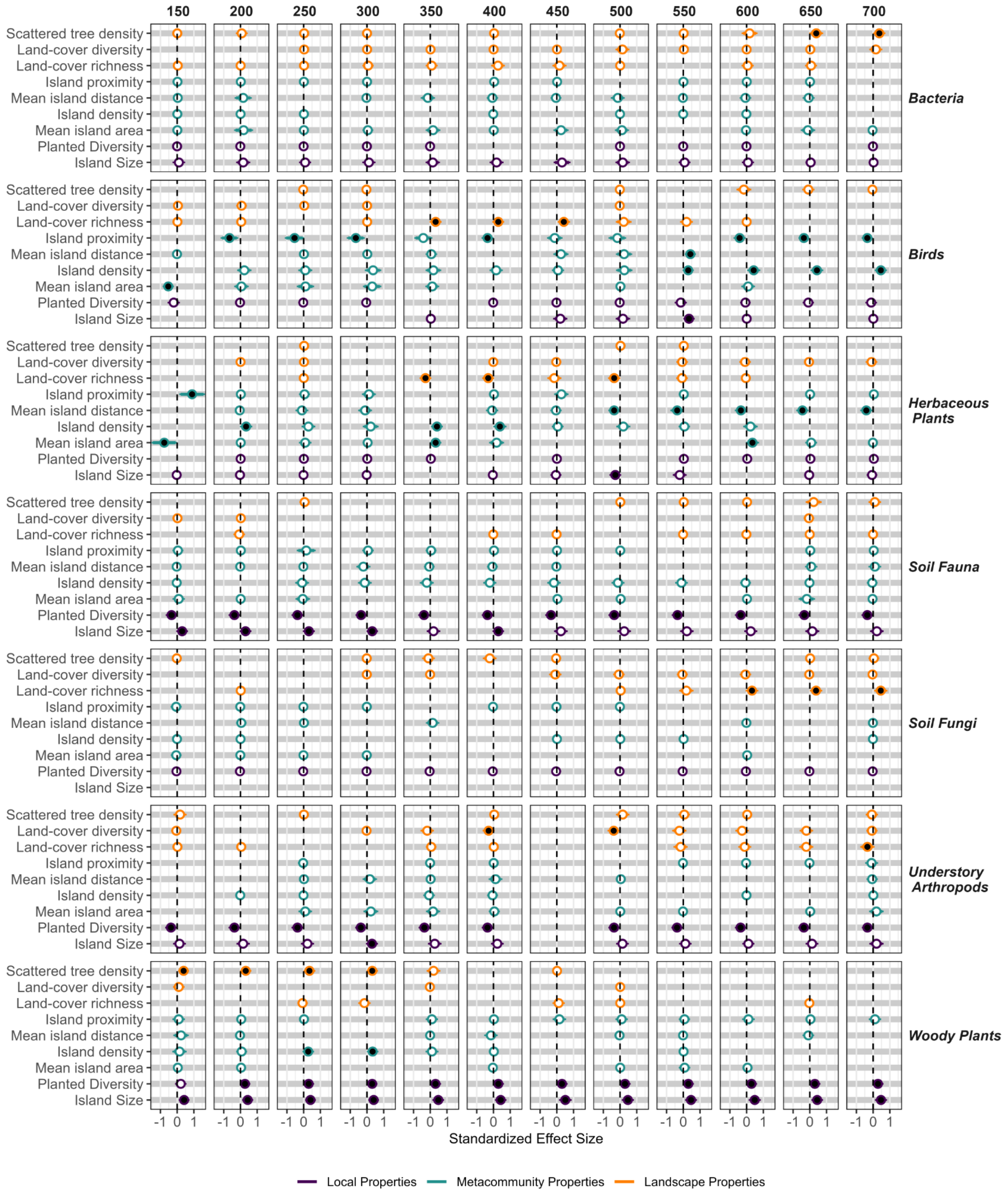


**Figure S8**. Comparison of the best-averaged multivariate models across spatial scales for each taxon, with panel headers identifying the spatial extent. Points with black fill indicate significant effects (p-values <= 0.05), while those with white fill represent non-significant effects (p-values

> 0.05). Effect sizes from the multivariate models are standardized, allowing for direct comparison across taxa and spatial scales.

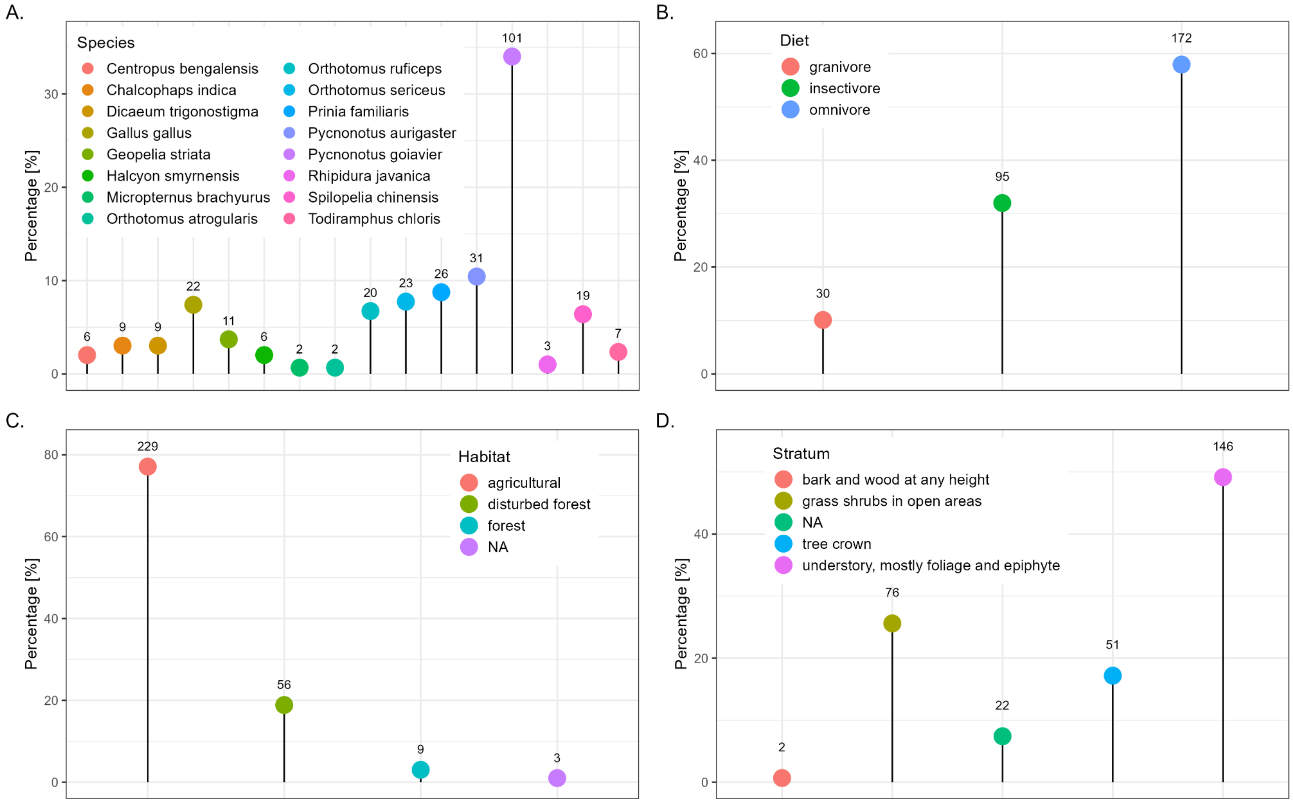


**Figure S9.** Summary of a) bird species composition, b) diet, c) preferred habitat, and d) habitat stratum as a percentage of the total. Counts are shown above each point.
